# *Entropy Fusion DNA:* Alignment-Free Gene Fusion Detection through Entropy and Mutual Information Descriptors

**DOI:** 10.64898/2026.05.27.728176

**Authors:** Gerardo Benevento, Delfina Malandrino, Alessia Ture, Rocco Zaccagnino

## Abstract

Gene fusions are clinically relevant genomic alterations and key cancer biomarkers. Their computational detection remains dominated by alignment-based pipelines, whose reliance on read mapping, reference annotations, and heuristic filtering makes them sensitive to mapping ambiguities, annotation incompleteness, repetitive regions, and false positives. Recent machine learning (ML) strategies aim to learn fusion-related patterns directly from sequencing data, but their adoption is still limited by dataset-specific biases, synthetic data artifacts, class imbalance, and representations that may overlook the structural organization of biological sequences.

Theoretical and statistical sequence descriptors remain underexplored as efficient tools for capturing informative structural signals in biological reads. In this work, we investigate whether fusion-related information can be inferred directly from the statistical organization of DNA sequences. Each sequence is encoded into a compact, interpretable, and alignment-free feature space combining Shannon and Rényi *entropy*, lagged and base-resolved *mutual information*, GC content, and rarefied *k*-mer richness descriptors. Our goal is to assess whether these information-theoretic features encode discriminative sequence signatures associated with fusion events.

For discriminating fusion-derived from non-fusion sequences, nested cross-validation selected K-nearest neighbors as the most effective classifier, achieving strong held-out performance on the balanced benchmark (AUROC = 0.892, AUPRC = 0.865). The same representation was then evaluated on fusion-positive samples for fusion partner prediction and breakpoint localization, achieving strong top-*k* partner identification accuracy and stable breakpoint regression performance. Moreover, a two-stage strategy in which the binary classifier first filters candidate reads further improved partner prediction, suggesting its use as an enrichment step for downstream fusion characterization. Although performance decreased under repeated fusion-pair-disjoint evaluation, it remained clearly above random expectation, supporting the transferability of the proposed descriptors to unseen fusion pairs. Breakpoint-centered validation further revealed increased local sequence complexity, altered short-range dependency structure, and modest but significant microhomology enrichment around fusion regions.

Such findings support an interpretable alignment-free framework where information-theoretic features provide predictive and biologically informative signals for gene fusion analysis.

The framework is available at: https://github.com/FLaTNNBio/EntropyFusionDNA

**Graphical Abstract:** 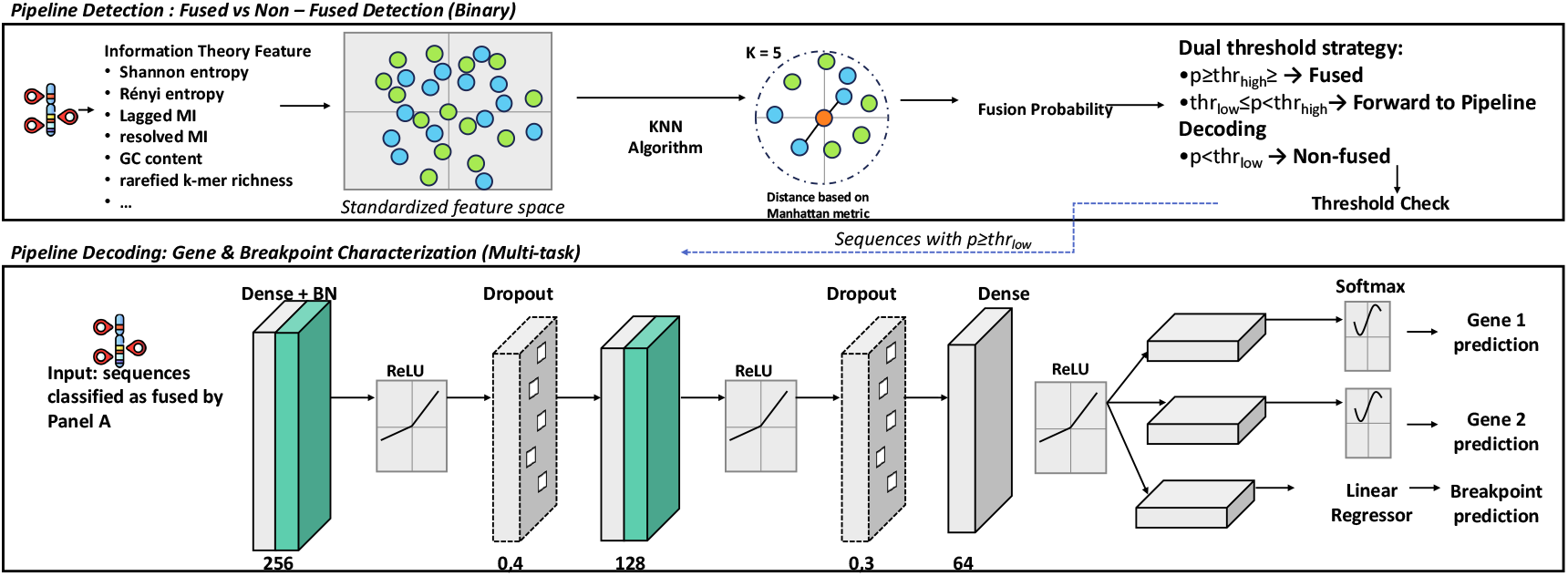

**Highlights:**

- Alignment-free information-theoretic DNA descriptors detect gene fusions.
- Resolved mutual-information features provide the strongest predictive signal.
- Two-stage screening enriches partner-gene prediction and breakpoint analysis.

## 1. Introduction

Recent advances in high-throughput sequencing technologies have transformed genomics by enabling the generation of data at unprecedented scale and resolution. In cancer research, these developments have improved disease understanding, diagnosis, and therapeutic decision-making through the systematic identification of mutations, structural rearrangements, and other clinically relevant genomic alterations. Among them, *gene fusion events* have gained particular attention, as they may function as oncogenic drivers, prognostic biomarkers, and actionable therapeutic targets across multiple tumour types. Gene fusions arise when distinct genomic loci are aberrantly joined through mechanisms such as chromosomal translocations, inversions, deletions, duplications, or transcriptional read-through events Mertens, Johansson, Fioretos and Mitelman (2015); Latysheva and Babu (2016). These events generate hybrid genes composed of portions of two or more original genes Annala, Parker, Zhang and Nykter (2013), leading to *chimeric* transcripts and, in many cases, fusion proteins with altered regulatory or functional properties. In humans, gene fusions are widely recognized as key contributors to tumour initiation, progression, and therapeutic response Liu, Nagasaka, Atz, Solca and Müllauer (2025). Clinically established examples, including *BCR-ABL1* and *EML4-ALK*, highlight their relevance as diagnostic biomarkers and therapeutic targets, making their reliable detection a central task in computational cancer genomics Hoskins, Vella, Reeser et al. (2025).

Most current fusion detection pipelines are alignment-based and analyze RNA-seq or whole-genome sequencing data through multistage workflows that integrate read mapping, discordant-read and split-read analysis, transcript assembly, and extensive heuristic filtering (McPherson, Hormozdiari, Zayed, Giuliany, Ha, Sun, Griffith, Heravi Moussavi, Senz, Melnyk et al., 2011; Kim and Salzberg, 2011). These approaches have had a major practical impact, but their performance can be affected by mapping ambiguities, local sequence repetitiveness, GC-content and coverage biases, incomplete reference genomes or transcript annotations, and the specific design of post-processing filters. Moreover, different fusion callers often show substantial disagreement in the set of reported events, so high-confidence predictions typically require consensus across multiple tools, followed by manual inspection.

More recent ML-based methods, such as *FusionAI*, have shown that DNA sequence windows surrounding known breakpoints contain predictive information. However, these approaches remain largely event-centric, as they typically rely on curated breakpoint-centered candidates as input (Kim, Tan, Liu, Yang and Zhou, 2021). This leaves open a complementary question: whether fusion-related signals can be inferred directly from the statistical organization of DNA sequences, independently of read-level alignment evidence. Indeed, fusion formation is not random with respect to genomic context. Breakpoints tend to occur in regions shaped by chromatin organization, replication timing, GC content, repetitive elements, and short sequence homologies (Hulke, Siefert, Sansam and Koren, 2019; Nambiar and Raghavan, 2011). These observations suggest that fusion-associated segments may carry detectable sequence-level signatures, and that such signatures could be captured through compact descriptors of sequence organization. In this perspective, an alignment-free and sequence-centric representation is attractive not only as a predictive strategy, but also as an interpretable framework for investigating the structural properties associated with gene fusion events.

In this work, we represent each DNA segment as a compact set of information theoretic and compositional descriptors, specifically, we model DNA as a symbolic process over the alphabet {***A, C, G, T***} and extract features including Shannon and Rényi entropy, lagged mutual information, base resolved mutual information, GC content, and rarefied *k* mer richness measures. Together, these descriptors summarize base composition, short range dependency structure, and higher order sequence organization without requiring explicit alignment to a reference genome. Our goal is not only to test whether such a representation can support fusion related prediction, but also to determine which components of sequence organization contribute most strongly to that signal.

We organize the analysis into four connected steps: *(i)* we address the primary task of distinguishing fused from non-fused sequences and show, through nested cross-validation, that this compact feature space supports strong predictive performance; among the models evaluated, K-nearest neighbors consistently emerged as the best classifier, outperforming both linear baselines and a feed-forward neural network, which indicates that fusion-related signal in this representation is captured more effectively by local, non-parametric decision structure than by simple global decision boundaries; *(ii)* we extend the framework to downstream tasks restricted to positive samples, namely fusion partner-gene prediction and breakpoint localization; *(iii)* we investigate the origin of the predictive signal through permutation importance and feature-family ablation analyses, showing that nucleotide-resolved mutual-information descriptors carry most of the discriminative information, whereas global compositional summaries contribute more modestly; and *(iv)* we assess robustness under stricter fusion-pair-disjoint evaluation and perform breakpoint-centered internal validation to test whether the learned signal is not only predictive, but also biologically coherent. Overall, our results support a concrete alignment-free strategy for gene-fusion sequence analysis and point to short-range nucleotide dependency patterns as a particularly informative direction for future method development.

## 2. Related Work

### 2.1. Gene fusions and computational approaches for their analysis

Gene fusions are a well-established class of oncogenic alterations across both haematological malignancies and solid tumours, where they can act as diagnostic markers, prognostic indicators, and therapeutic targets (Mertens et al., 2015; Tomlins, Rhodes, Perner, Dhanasekaran, Mehra, Sun, Varambally, Cao, Tchinda, Kuefer et al., 2005). Canonical examples include *BCR-ABL1* in chronic myeloid leukaemia and *ETV6-RUNX1* in childhood acute lymphoblastic leukaemia, although large sequencing studies have shown that fusion events are widespread across tumour types and contribute substantially to tumour heterogeneity and molecular stratification (Chandran, Geetha, Sakthivel, Aswathy, Gopinath, Raj, Priya, Nair and Sreedharan, 2019; De Braekeleer, Douet-Guilbert, Guardiola, Rowe, Mustjoki, Zamecnikova, Al Bahar, Jaramillo, Berthou, Bown et al., 2013; Lilljebjörn, Henningsson, Hyrenius-Wittsten, Olsson, Orsmark-Pietras, Von Palffy, Askmyr, Rissler, Schrappe, Cario et al., 2016; Zelent, Greaves and Enver, 2004; Annala et al., 2013; Gatalica, Xiu, Swensen and Vranic, 2019). Their formation can arise through multiple genomic mechanisms, including balanced translocations, complex rearrangements, template-switching processes, and catastrophic chromosome shattering events, and is influenced by broader genomic context such as chromatin organisation, genome instability, replication-related processes, and three-dimensional chromosome architecture (Weischenfeldt, Symmons, Spitz and Korbel, 2013; Nambiar and Raghavan, 2011; Zhang, Spektor, Cornils, Francis, Jackson, Liu, Meyerson and Pellman, 2015; Kaiser and Semple, 2018; Hnisz, Schuijers, Lin, Weintraub, Abraham, Lee, Bradner and Young, 2015; Hulke et al., 2019; Drier, Lawrence, Carter, Stewart, Gabriel, Lander, Meyerson, Beroukhim and Getz, 2013). From a computational perspective, most practical fusion discovery pipelines are alignment based and operate primarily on RNA-seq data. Tools such as *deFuse, TopHat-Fusion, STAR-Fusion*, and *FusionCatcher* detect candidate fusion transcripts by combining splice-aware alignment, discordant and split-read evidence, transcript reconstruction, and extensive post-processing heuristics (McPherson et al., 2011; Kim and Salzberg, 2011; Haas, Dobin, Stransky, Li, Yang, Tickle, Bankapur, Ganote, Doak, Pochet et al., 2017; Nicorici, Şatalan, Edgren, Kangaspeska, Murumägi, Kallioniemi, Virtanen and Kilkku, 2014). These methods have become the dominant paradigm for fusion detection, but their performance depends strongly on read mappability, repetitive sequence content, sequencing depth, GC bias, and the quality of reference genome and transcript annotation. Large benchmarking studies have shown that fusion callers often disagree substantially, reflecting different trade-offs between sensitivity, precision, and filtering strategy (Haas, Dobin, Li, Stransky, Pochet and Regev, 2019).

Alongside these alignment based methods, machine-learning approaches have increasingly been proposed to score or characterize fusion *events. FusionAI*, for example, applies deep learning to breakpoint-centred DNA windows in order to learn sequence patterns associated with plausible fusion breakpoints (Kim et al., 2021). *DEEPrior* instead addresses the prioritisation of already detected gene fusions by estimating their oncogenic relevance downstream of the calling step (Lovino, Ciaburri, Urgese, Di Cataldo and Ficarra, 2020). These studies support the idea that useful fusion-related signal is embedded in the local sequence context. However, they remain largely event centric, in the sense that they typically assume candidate partner genes or breakpoint-centred regions are already available. This leaves open a different question, namely how far compact handcrafted descriptors of sequence organization can support fusion-related inference in a more general alignment free and segment centric setting.

### 2.2. Information-theoretic and *k*-mer-based representations of biological sequences

Information-theoretic descriptors and alignment free oligonucleotide statistics have long been used to characterize biological sequences. Entropy based, complexity based, and compression inspired analyses have been applied to quantify sequence heterogeneity, identify low-complexity regions, segment genomes into compositionally distinct domains, and distinguish different classes of biological sequence organization (Wootton and Federhen, 1993; Schmitt and Herzel, 1997; Bernaola-Galván, Grosse, Carpena, Oliver, Román-Roldán and Stanley, 2000; Grosse, Bernaola-Galván, Carpena, Román-Roldán, Oliver and Stanley, 2002; Koslicki, 2011; Damaševičius, 2010; Vinga, 2014). In these approaches, DNA is treated as a symbolic stochastic process, and deviations from randomness or independence are interpreted as signatures of biological structure.

Among these descriptors, mutual information is especially relevant because it captures dependency structure beyond single-nucleotide composition. Mutual-information-based analyses have been used to quantify short-range correlations, periodicities, and context-dependent organization in DNA sequences, revealing informative structure that cannot be recovered from marginal base frequencies alone (Kugiumtzis and Provata, 2004; Damaševičius, 2010). The average mutual information profile has also been proposed as a genomic signature, supporting the broader idea that dependency patterns can serve as compact but informative sequence fingerprints (Bauer, Schuster and Sayood, 2008). More recently, resolved formulations of mutual information have been introduced as structural fingerprints of biomolecular sequences for interpretable machine learning, making it possible to decompose dependency patterns by residue or nucleotide identity and thereby improve interpretability (Bohnsack, Kaden, Abel, Saralajew and Villmann, 2021). This direction is particularly relevant in genomic settings where subtle local sequence constraints may carry biological significance. In parallel, *k*-mer composition and related alignment free representations have become standard tools in comparative genomics and sequence analysis (Vinga and Almeida, 2003; Bonham-Carter, Steele and Bastola, 2014). Early work on dinucleotide and oligonucleotide usage established the concept of genome signatures, showing that genomes display stable and non-random compositional biases that can be exploited for classification and comparison (Kariin and Burge, 1995). Extensions to longer word statistics and background adjusted alignment free dissimilarity measures have supported applications ranging from phylogeny and metagenomics to transfer detection and anomaly analysis (Qi, Wang and Hao, 2004; Tang, Lu and Sun, 2018). Together, information-theoretic descriptors and *k*-mer-based statistics provide a principled way to describe the statistical texture of DNA without explicit reference alignment.

### 2.3. Positioning of the present work

The present study lies at the intersection of gene fusion analysis and alignment free sequence modelling. On the one hand, it is motivated by growing evidence that fusion breakpoints and rearrangements are shaped by non-random genomic context, including local sequence properties, DNA breakage mechanisms, and broader structural features of the genome (Nambiar and Raghavan, 2011; Drier et al., 2013; Kaiser and Semple, 2018; Hulke et al., 2019). On the other hand, it builds on the long tradition of representing biological sequences through compact and interpretable information-theoretic and oligonucleotide-based descriptors (Vinga, 2014; Bonham-Carter et al., 2014; Bauer et al., 2008; Bohnsack et al., 2021). Unlike alignment based fusion callers such as *STAR-Fusion* and *FusionCatcher*, our framework does not rely on read mapping, split-read support, or transcript reconstruction. Unlike event centric learning methods such as *FusionAI* and *DEEPrior*, it does not assume that candidate fusion events have already been defined. Instead, we adopt a segment centric formulation in which each DNA sequence is represented by a small set of global and local handcrafted descriptors, and we investigate how far this interpretable feature space can support both prediction and biological interpretation in the context of gene fusions. In this sense, the present work is not only a modelling exercise, but also an attempt to identify which specific statistical properties of DNA sequence organization are most strongly associated with fusion-related genomic regions.

## 3. Materials and Methods

The methodological framework developed in this study combines sequence representation, predictive modeling, and internal biological validation within a unified alignment-free setting. In the following subsections, we describe the construction of the fused and non-fused dataset, the extraction of handcrafted information-theoretic descriptors, and the feature representation used for model comparison. We then present the binary-classification benchmark and nested cross-validation strategy used to select the most suitable classifier, followed by the held-out evaluation of the final model, post hoc interpretability analyses, and the second-stage multitask framework for partner-gene and breakpoint prediction. The section concludes with robustness experiments, breakpoint-centered and microhomology-based internal validation, and comparison with FusionAI-like and alignment-based approaches.

### 3.1. Study design overview

The study was designed as a sequence-centric and alignment-free analysis of gene-fusion-associated DNA segments, with the dual aim of evaluating predictive performance and validating the biological relevance of compact information-theoretic sequence descriptors. The overall workflow consisted of five stages: (i) assembly of a dataset of fused and non-fused genomic sequences derived from curated fusion resources and matched controls; (ii) representation of each sequence through a compact feature set based on entropy, mutual information, GC content, and rarefied

*k*-mer richness statistics; (iii) evaluation of the primary binary-classification task between fused and non-fused sequences through comparison of multiple machine-learning models under nested cross-validation; (iv) extension of the framework to downstream tasks restricted to positive samples, namely prediction of fusion partner genes and breakpoint position; and (v) interpretability, robustness, and internal biological validation analyses aimed at determining whether the most informative features captured sequence properties that were not only predictive, but also biologically meaningful in the context of fusion formation.

Additional experiments were performed under more demanding evaluation settings, including fusion-pair-disjoint testing, FusionAI-like benchmarking, and comparison with alignment-based fusion callers.

### 3.2. Dataset construction

Positive samples were derived from curated gene-fusion events collected from *FusionGDB2* (Kim, Tan, Liu, Lee, Jung, Kumar and Zhou, 2022). The original positive dataset contained 116,126 fusion-associated sequence records, each represented by a DNA sequence, a binary class label, the two partner-gene identifiers, and the junction coordinate defining the fusion breakpoint within the sequence. Before constructing the final dataset, positive records were deduplicated using the combination of first partner gene, second partner gene, junction point, and sequence as the duplicate key. This procedure removed 34,753 duplicate entries, retaining 81,373 unique positive fusion instances. Negative samples were generated as artificial chimeric controls using transcript sequences from the GENCODE v19/GRCh37 protein-coding reference. For each retained positive instance, one negative sequence was constructed in order to obtain a balanced dataset. Each negative sample was generated by selecting two genes from the pool of genes observed among the positive fusion instances, with sampling probabilities proportional to their marginal frequencies as first or second fusion partners in the positive set. This strategy was adopted to reduce trivial differences in gene composition between the two classes. Candidate gene pairs corresponding to curated positive fusion pairs were excluded, as were their reverse configurations and self-pairs. After selecting the two genes, one transcript-derived segment was randomly extracted from each gene according to the reference annotation, with segment lengths sampled between 60 and 5000 nucleotides. The artificial negative sequence was then obtained by concatenating the segment sampled from the first gene with the segment sampled from the second gene. The junction point was defined as the length of the first segment, thereby marking the exact concatenation site between the two transcript-derived regions. These synthetic chimeric sequences were labelled as non-fused controls (*label* = 0), since they did not correspond to curated or observed gene-fusion events. Both positive and negative samples were stored using the same representation, including the nucleotide sequence, binary label, first partner gene, second partner gene, and junction point. This design ensured that the classification task could not rely on class-specific placeholder values, missing metadata fields, or structural differences in the input format. The final dataset contained 162,746 sequences, comprising 81,373 positive fusion samples and 81,373 artificial chimeric negative controls, and was used as the starting point for subsequent preprocessing, feature extraction, model selection, and evaluation procedures.

### 3.3. Class balancing and data partitioning

The dataset was constructed to be balanced at the class level. After positive deduplication, all 81,373 unique positive fusion instances were retained, and one artificial chimeric negative control was generated for each positive instance. This create a final benchmark of 162,746 sequences with an equal number of positive and negative samples. Data partitioning was performed in a stratified manner so that class proportions were preserved across folds and held-out subsets. The same principle was applied both in the inner and outer loops of nested cross-validation during model selection and hyperparameter optimization. Whenever explicit train, validation, and test splits were required, these were generated under the same stratified scheme, while random sampling, artificial negative generation, and data partitioning were performed using fixed random seeds to ensure reproducibility.

### 3.4. Information theoretic and compositional feature extraction

Each DNA sequence was represented as a symbolic string over the alphabet 𝒜 = {*A, C, G, T* }. Before feature extraction, sequences were cleaned by removing characters outside this alphabet, so that all descriptors were computed on the filtered nucleotide string only. From each sequence, we extracted a compact alignment-free feature space designed to summarize statistical structure at the levels of base composition, short-range dependency, and oligonucleotide diversity.

Let *s*_1_, …, *s*_*L*_ denote a cleaned DNA sequence of length *L*. For each nucleotide *x ∈* 𝒜, let 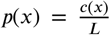 be its empirical frequency, where *c*(*x*) is the number of occurrences of *x* in the sequence.

#### Shannon entropy

Global nucleotide uncertainty was quantified through Shannon entropy:

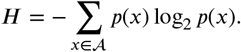

#### Rényi entropy

To complement Shannon entropy with a measure more sensitive to concentration effects, we also computed Rényi entropy of order *α*:

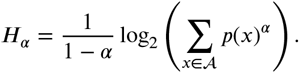

In the present study, the retained Rényi descriptor corresponded to order *α* = 2.

#### Lagged mutual information

To characterize short-range dependencies between nucleotides, we computed mutual information at lag *τ*:

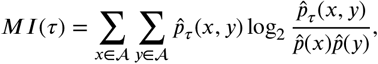

Where 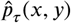 denotes the smoothed empirical joint probability of observing nucleotide *x* at position *i* and nucleotide *y* at position *i* +*τ*, and 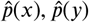are the corresponding smoothed marginals. In the implementation, these probabilities were estimated using add-*α* smoothing with *α* = 0.5:

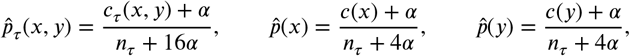

where *c*_*τ*_ (*x, y*) is the number of valid (*x, y*) pairs at lag *τ, c*(*x*) and *c*(*y*) are the associated marginal counts, and *n*_*τ*_ is the total number of valid nucleotide pairs at that lag. A Miller–Madow bias correction was additionally applied to the global mutual information estimate. In the final representation, lagged mutual information was evaluated for *τ* = 1, …, 5.

#### Resolved mutual information

To obtain a more detailed view of local dependency structure, we also computed nucleotide-resolved mutual information descriptors. For a given lag *τ* and nucleotide *x ∈* 𝒜, the resolved mutual information was defined as

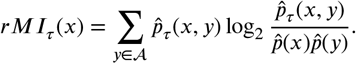

Thus, *rMI*_*τ*_ (*x*) measures the contribution of nucleotide *x* to the lagged mutual information at lag *τ*, and by construction

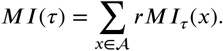

Resolved mutual information descriptors were computed for each nucleotide *x ∈* {*A, C, G, T* } and each lag *τ* = 1, …, 5, resulting in 20 variables in the final representation. No additional Miller–Madow correction was applied to the nucleotide-resolved terms.

#### GC content

Global base composition was further summarized through GC content:

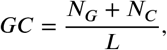

where *NG* and *NC* are the counts of nucleotides G and C in the sequence.

#### k-mer richness features

To quantify higher-order compositional diversity, we computed rarefied unique *k*-mer richness ratios for *k* = 3, 4, 5. For a sequence of length *L*, the total number of overlapping *k*-mers is *L* − *k* + 1. Let *mk* = min(1000, *L*−*k*+1). We then sampled *mk k*-mer positions without replacement and defined the rarefied richness ratio as

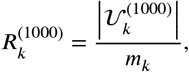

where 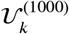 is the set of distinct *k*-mers observed in the rarefied sample. These descriptors summarize oligonucleotide d*k*iversity while reducing the effects of sequence-length variability and sampling noise.

### 3.5. Final features representation

The final feature representation used in the main benchmark experiments consisted of 31 handcrafted variables drawn from five descriptor families, namely entropy-based descriptors, lagged mutual information, resolved mutual information, GC content, and rarefied *k*-mer richness ratios. More specifically, the final set comprised two global entropy measures, five lagged mutual-information descriptors for *d* = 1, …, 5, twenty nucleotide-resolved mutual-information variables obtained across four nucleotides and five lags, one GC-content descriptor, and three rarefied unique *k*-mer richness ratios for *k* = 3, 4, 5 (Table 1). Raw sequence-length information, together with the non-rarefied unique *k*-mer ratio descriptors, was excluded from the final benchmark representation, since our aim was to retain a compact feature space that emphasized interpretable compositional and dependency-based properties while reducing redundancy and trivial length-related effects. By using the same 31-feature input space for all candidate classifiers, we ensured that model comparison reflected differences in learning behavior rather than differences in available input information. At the same time, because a central objective of the study was not only predictive performance but also interpretability, this final 31-feature representation provided the common basis for all subsequent analyses, including feature-importance estimation, feature-family ablation, robustness assessment, and internal biological validation.

**Table 1.** Composition of the final 31 feature representation used in the main benchmark experiments.

| Feature family | Number of features | Included descriptors |
| --- | --- | --- |
| Entropy based descriptors | 2 | Shannon entropy; Rényi entropy ($\alpha = 2$) |
| Lagged mutual information | 5 | $MI_{\tau=1}, MI_{\tau=2}, MI_{\tau=3}, MI_{\tau=4}, MI_{\tau=5}$ |
| Resolved mutual information | 20 | $rMI_{A,\tau=1\dots5}, rMI_{C,\tau=1\dots5}, rMI_{G,\tau=1\dots5}, rMI_{T,\tau=1\dots5}$ |
| GC content | 1 | GC content |
| Rarefied $k$-mer richness | 3 | rarefied unique 3-mer ratio; rarefied unique 4-mer ratio; rarefied unique 5-mer ratio |
| Total | 31 | — |

### 3.6. Primary task: fused versus non fused classification

The primary predictive task in this study was binary classification of DNA segments as fused or non fused. To ensure a controlled comparison across learning strategies, all candidate models were trained on the same balanced benchmark dataset of 162,962 sequences and on the same 31 feature representation described above, so that differences in performance could be attributed to the learning algorithm rather than to differences in input information.

We compared four families of classifiers, namely Logistic Regression, Linear Support Vector Classification (`LinearSVC`), K nearest neighbors (KNN), and a feed forward neural network baseline. Logistic Regression and LinearSVC were included as linear baselines, since they provide simple decision boundaries and therefore offer a useful reference for evaluating whether the handcrafted representation is linearly separable, whereas KNN was included as a non parametric local classifier and the neural model as a nonlinear parametric baseline capable of learning more complex transformations of the same input space.

For the classical models, preprocessing and classification were implemented through scikit learn pipelines in which `StandardScaler` was applied before the classifier. Hyperparameter optimization was then carried out over model specific grids, with the regularization strength varied over *C ∈* {0.01, 0.1, 1, 10} for Logistic Regression and LinearSVC, while for KNN we explored the number of neighbors *k ∈* {1, 3, 5, 11, 15, 31, 51, 101}, the weighting scheme {`uniform, distance`}, and the Minkowski distance parameter *p ∈* {1, 2}, corresponding to Manhattan and Euclidean distance, respectively.

The neural network baseline was trained on the same standardized 31 feature input and was included to test whether a nonlinear parametric model could exploit structure in the handcrafted representation beyond what was accessible to linear or neighborhood based classifiers. The details of its architecture, hyperparameter tuning, and training procedure are described in the following subsection.

### 3.7. Model selection by nested cross validation

Model selection and hyperparameter optimization were performed by nested cross validation so as to reduce optimistic bias and obtain a more reliable estimate of generalization performance under model tuning. For the classical models, we adopted a 5 fold stratified outer cross validation loop together with a 3 fold stratified inner loop, so that hyperparameters could be optimized independently within each outer training partition while preserving class balance throughout the procedure. In each outer fold, candidate hyperparameter settings were compared on the corresponding inner folds using AUROC as the selection criterion, after which the best estimator was retrained on the full outer training split and evaluated on the associated outer validation fold. The neural network baseline was evaluated under the same general nested cross validation structure, again using 5 stratified outer folds and 3 stratified inner folds, although its training procedure required an explicit separation between preprocessing, hyperparameter selection, and early stopping. Accordingly, within each outer fold a `StandardScaler` was fitted on the outer training data only and then applied unchanged to both the inner validation subsets and the final outer validation subset. Hyperparameter selection in the inner loop considered 16 configurations obtained from the Cartesian product of learning rate {10^−3^, 3 × 10^−4^}, first layer dropout {0.4, 0.2}, second layer dropout {0.3, 0.1}, and weight decay {0, 10^−4^}. For each inner split, the model was trained with early stopping based on validation loss, while the mean inner fold AUROC was used to select the best hyperparameter configuration. Once selected, the corresponding model was retrained on the full outer training split and then evaluated on the associated outer validation fold. Performance was summarized across outer folds using AUROC, AUPRC, accuracy, and F1 score, and for each metric we report the mean across outer folds together with an approximate 95% confidence interval computed from the standard error of the fold wise scores. The classifier achieving the best nested cross validation performance was then chosen as the main model for subsequent binary classification analyses, whereas the comparative performance estimates obtained under this protocol are reported in Table 2 and discussed in the Results section.

**Table 2.** Performance of the candidate classifiers in 5-fold outer nested cross-validation on the balanced 31-feature benchmark dataset. Hyperparameters were selected independently within each outer-training split by 3-fold inner cross-validation. Reported values are mean scores across outer folds with approximate 95% confidence intervals. Best values are shown in bold.

|  |  |  |  |  |
| --- | --- | --- | --- | --- |
| Logistic Regression | 0.665 [0.663, 0.668] | 0.627 [0.624, 0.630] | 0.620 [0.617, 0.624] | 0.659 [0.656, 0.662] |
| LinearSVC | 0.665 [0.663, 0.668] | 0.628 [0.625, 0.630] | 0.620 [0.617, 0.623] | 0.661 [0.658, 0.664] |
| Feed-forward neural network | 0.849 [0.847, 0.851] | 0.845 [0.842, 0.849] | 0.767 [0.764, 0.769] | 0.772 [0.770, 0.775] |
| K-nearest neighbors | **0.905 [0.903, 0.907]** | **0.879 [0.877, 0.881]** | **0.808 [0.805, 0.810]** | **0.826 [0.824, 0.828]** |

### 3.8. Final KNN model and held out evaluation

Following model selection by nested cross validation, K nearest neighbors was selected as the reference classifier for fused versus non fused discrimination, and all final held out analyses were therefore carried out using a KNN model with *k* = 5, distance based weighting, and Minkowski distance with *p* = 1, which corresponds to a Manhattan distance metric. For these analyses, the balanced dataset was partitioned into training, validation, and test subsets, after which continuous input variables were standardized by centering each feature to zero mean and scaling it to unit variance. To avoid information leakage, feature standardization was fitted on the training set only and then applied unchanged to the validation and test sets. Class probabilities were obtained directly from the posterior estimates returned by the KNN classifier. Because the binary classifier was used in two settings with different priorities, decision thresholds were defined explicitly on the validation set rather than fixing a single default cutoff. This allowed us to distinguish between an operating point optimized for balanced standalone classification performance and a more permissive operating point designed to retain candidate fused sequences for downstream analysis. For thise reason, two decision thresholds were then defined on the validation set, and each was used for a distinct analytical purpose:

i. thr_*high*_, selected by maximizing the validation F1 score, was used as the main operating threshold for the standalone fused versus non fused classification task, since it provided the best balance between precision and recall under the primary binary setting;
ii. thr_*low*_, selected as the highest threshold achieving validation recall of at least 0.90, was used as a more permissive screening threshold in the downstream two stage analysis, where preserving potentially fused candidates was more important than maximizing first stage precision.

Performance on the held out test set was summarized using AUROC, AUPRC, accuracy, precision, recall, and F1 score, while confidence intervals were estimated by bootstrap resampling of the test set in order to quantify the stability of the reported metrics.

### 3.9. Post hoc feature importance and feature family ablation

After selecting KNN as the primary classifier for the fused versus non fused task, we performed two complementary post hoc analyses in order to characterize the contribution of individual descriptors and descriptor families within the final 31 feature representation. In particular, the first analysis was designed to quantify the importance of single variables within the trained classifier, whereas the second was intended to determine how much predictive performance depended on broader groups of related descriptors.

i. **Permutation based feature importance**. Feature importance was assessed on the validation set of the first stage binary classifier by independently permuting one feature at a time while keeping all remaining variables unchanged, after which predictive performance was recomputed and compared with the unpermuted baseline. Importance was quantified as the performance decrease induced by permutation and was measured in terms of AUROC, AUPRC, accuracy, F1 score, precision, and recall. To reduce Monte Carlo variability, each feature was permuted repeatedly and the mean degradation across repetitions was recorded, while for computational tractability the analysis could be performed on a fixed size random subsample of the validation set.
ii. **Feature family ablation**. We then carried out a feature family ablation analysis using the same KNN architecture, so as to assess whether predictive performance depended mainly on individual descriptors or on broader classes of sequence statistics. This analysis was performed in two complementary modes, namely a leave one group out setting in which one descriptor family was removed from the full representation and the classifier was retrained from scratch, and a group only setting in which the classifier was trained using only one descriptor family at a time. The feature families considered in this analysis corresponded to the five descriptor classes retained in the final representation, namely entropy based descriptors, lagged mutual information, resolved mutual information, GC content, and rarefied unique *k*-mer richness measures. Together, these analyses were intended to determine whether performance was driven primarily by global compositional summaries or by local dependency patterns, especially those captured by mutual-information-based descriptors.

### 3.10. Secondary tasks on positive samples: partner-gene and breakpoint prediction

To determine whether the same 31-feature representation could support a finer characterization of fusion-positive sequences, we introduced a second-stage multitask model, here referred to as the Decoding stage, and restricted its training and evaluation to positive samples only. This analysis moved beyond binary fused-versus-non-fused discrimination and addressed three related objectives simultaneously: prediction of the first partner-gene label (*gene1*), prediction of the second partner-gene label (*gene2*), and prediction of the breakpoint position.

The fused-only subset comprised 81,481 positive samples and, for each sequence, we retained the same 31 information-theoretic and compositional descriptors used in the primary binary-classification task, together with the associated annotations *gene1, gene2*, the raw breakpoint coordinate (`junction_point`), and sequence length. To make breakpoint prediction comparable across sequences of different length, the breakpoint coordinate was normalized as rel_bp = junction_point/seq_len, and the resulting values were clipped to the interval [0, 1].

The union of all genes appearing as either *gene1* or *gene2* in the fused subset defined the label vocabulary for the two partner-prediction tasks, corresponding to 9,866 unique genes overall. To ensure consistency with the first-stage Detection model, the fused-only samples inherited the same master train, validation, and test partition used for the binary task, after restricting each split to positive instances only. This resulted in 55,407 training samples, 9,778 validation samples, and 16,296 test samples. Input features were standardized by fitting the scaling transformation on the training subset only and then applying the same transformation unchanged to validation and test data.

The Decoding model was selected by validation-based tuning over a small grid of candidate multitask feed-forward architectures. Candidate shared backbones with hidden-layer sizes 128, 64, 64, 256, 128, 128, and 384, 192, 128 were combined with dropout pairs (0.4, 0.3) and (0.2, 0.1), while the learning rate was fixed at 10^−3^. Model selection was based on validation loss, and the best-performing configuration was retained as the final Decoding model.

The final multitask network consisted of a shared backbone with hidden layers of sizes 384, 192, and 128, followed by three task-specific heads: one linear classifier for *gene1*, one linear classifier for *gene2*, and one linear regression head for breakpoint prediction. Training was performed with the Adam optimizer, a batch size of 256, and early stopping with patience 10. The multitask objective combined the two cross-entropy losses for partner-gene prediction with the mean squared error loss for breakpoint regression:

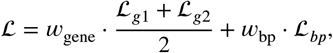

with 𝓌_gene_ = 1.0 and 𝓌_bp_ = 0.5.

Partner-gene prediction was evaluated using top-1, top-3, and top-5 accuracy for both *gene1* and *gene2*, whereas breakpoint prediction was summarized using mean absolute error and mean squared error on the normalized breakpoint coordinate. The Decoding stage was evaluated in two complementary settings: an *Oracle* setting, in which the multitask model was applied directly to fused test samples, and a combined *Detection+Decoding* setting, in which Decoding was applied only to fused test instances that also received sufficiently high fusion probability from the first-stage KNN Detection model under either thr_*low*_ or thr_*high*_.

### 3.11. Robustness under fusion pair disjoint evaluation

To assess robustness under a stricter generalization setting, we repeated the primary fused versus non fused classification task under a fusion pair disjoint protocol. In this setting, training and test samples were separated so that fusion pair identities present in the test partition were completely absent from the training partition. As a result, the model could not benefit from having previously seen the same fusion partner combination during training.

For example, if a sequence involving the fusion pair *ETV6-RUNX1* was assigned to the test set, no other instance involving the same *ETV6-RUNX1* pair was allowed to appear in the corresponding training set. This makes the task substantially more demanding than an ordinary random split, because the classifier must rely on transferable sequence level properties rather than on pair specific similarity or repeated fusion identities. Robustness was evaluated on the balanced 31 feature dataset using repeated grouped splits with zero train test overlap in fusion pair identity. Both KNN and the feed forward neural network were tested under this protocol. Performance was summarized across repeated splits using AUROC, AUPRC, accuracy, F1 score, precision, and recall. This protocol was designed to stress the models and to test whether the proposed feature space captures general properties of fusion associated sequences rather than signals tied to recurrent fusion pairs.

### 3.12. Breakpoint centered internal validation

To evaluate the biological coherence of the handcrafted representation, we performed an internal breakpoint-centered validation analysis. Fixed-length windows were extracted around annotated fusion breakpoints and compared with matched windows from non-fused control sequences positioned at the same relative location within the segment. Local information-theoretic descriptors were then recomputed within each window, thereby restricting the analysis to the immediate breakpoint neighborhood. This analysis was designed to determine whether the feature families that emerged as important in the global fused-versus-non-fused classification task also retained discriminatory power in local breakpoint-centered sequence contexts. Each descriptor was evaluated individually by univariate AUROC, while distributional separation between fusion and non-fusion windows was quantified using Cliff’s delta and the Mann–Whitney U test. Thus, the breakpoint-centered validation served as an internal biological check on whether the predictive signal identified at the whole-sequence level was also reflected in interpretable local properties of true fusion breakpoint regions.

### 3.13. Microhomology based internal validation

Because short stretches of sequence homology are a plausible mechanistic contributor to DNA breakage and rearrangement repair, we performed an additional internal validation based on breakpoint microhomology. For sequences with valid breakpoint annotations, microhomology length was quantified as the number of identical nucleotides shared across the breakpoint junction, that is, the maximal exact overlap between the two sides of the joined sequence. Fusion positive samples were then compared with matched non fused controls in order to test whether authentic fusion regions were enriched in local sequence contexts compatible with microhomology mediated rearrangement processes. The analysis was designed as a conservative matched comparison. Fusion positive samples were paired with non fused controls represented in the same sequence format, so that differences in microhomology distribution would reflect local breakpoint related sequence properties rather than trivial differences in sequence construction. Microhomology distributions were summarized in terms of mean and median overlap length, fraction of non zero events, univariate AUROC, Cliff’s delta, and Mann–Whitney testing. To relate this mechanistic property to the handcrafted representation used throughout the study, we also examined the association between breakpoint microhomology length and the global descriptors retained in the final 31 feature representation. In addition, we assessed the relationship between microhomology and the KNN fusion score assigned by the first stage classifier. These analyses were intended to determine whether part of the predictive signal captured by the handcrafted descriptors could be linked to a biologically plausible breakpoint property, while also testing whether microhomology alone could account for the observed classification performance. In this way, the microhomology analysis served as a complementary internal validation of biological plausibility rather than as an alternative predictive model.

### 3.14. Comparison with FusionAI like and alignment based baselines

To contextualize the proposed framework, we compared it with both sequence based and alignment based baselines. For *FusionAI like* evaluation, we constructed a matched 1:1 dataset consisting of 26,000 real fused sequences and 26,000 pseudo fusion negatives. Pseudo fusion negatives were generated by combining transcript derived sequences from genes not represented in the positive fusion set, and the same 31 handcrafted features were computed for all resulting sequences. Performance was then evaluated using a 70:30 train test split in order to mimic a candidate based sequence classification protocol. For comparison with alignment based fusion callers, we considered representative event level tools including *FusionCatcher* and *STAR Fusion*. Since these tools report fusion events rather than labels for pre extracted sequence instances, their outputs were mapped to the benchmark through an event to instance matching procedure. In this comparison, a sequence level instance was considered positive when it contributed to a reported fusion event under the adopted evaluation mapping. These comparisons were intended to position the proposed framework relative to established paradigms, while recognizing that event level callers and segment level classifiers operate under different assumptions.

## 4. Results

### 4.1. Nested cross-validation identifies K-nearest neighbors as the best classifier

The nested-cross-validation comparison revealed a clear and highly consistent hierarchy among the candidate classifiers, with K-nearest neighbors emerging as the strongest model on the balanced fused-versus-non-fused benchmark and with all performance estimates reported in Table 2. Because these values are based exclusively on outer-fold evaluation, after hyperparameter selection had been completed independently within each outer-training partition, they provide an approximately unbiased estimate of generalization performance under model selection and therefore offer the most reliable basis for comparing the alternative learning strategies considered here.

The most important result is that KNN separated fused from non-fused sequences substantially better than all competing models, reaching a mean AUROC of 0.905 (95% CI [0.903, 0.907]) and a mean AUPRC of 0.879 (95% CI [0.877, 0.881]), while also achieving the best threshold-dependent performance with a mean accuracy of 0.808 and a mean F1-score of 0.826. This advantage is not marginal but pronounced, and it immediately suggests that the discriminative structure induced by the handcrafted descriptors is organized primarily in terms of local neighborhood relationships rather than through a simple global separating surface. This interpretation is reinforced by the behavior of the two linear baselines, which performed much worse and remained nearly indistinguishable from one another. Logistic Regression reached a mean AUROC of 0.665 and a mean AUPRC of 0.627, whereas LinearSVC achieved a mean AUROC of 0.665 and a mean AUPRC of 0.628, with similarly modest values for accuracy and F1-score. Taken together, these results indicate that fused and non-fused sequences are not well separated by a linear decision boundary in the selected 31-dimensional feature space, even though the underlying descriptors are biologically interpretable and globally informative.

The feed-forward neural-network baseline occupied an intermediate position, and although it improved markedly over both linear models, thereby confirming the presence of nonlinear discriminative structure in the representation, it still remained clearly below KNN across all reported metrics. Across the five outer folds, the neural model reached a mean AUROC of 0.849 (95% CI [0.847, 0.851]) and a mean AUPRC of 0.845 (95% CI [0.842, 0.849]), together with a mean accuracy of 0.767 and a mean F1-score of 0.772. Thus, nonlinear modeling is clearly beneficial, yet the best performance is obtained not by a parametric transformation of the feature space but by a classifier that exploits local similarity structure directly. A further point supporting the robustness of this result is that hyperparameter selection for KNN was completely stable across outer folds. In every fold, the same configuration was selected, namely *k* = 5, distance-based neighbor weighting, and a Minkowski metric with *p* = 1, corresponding to Manhattan distance.

This stability indicates that the superiority of KNN was not the consequence of fold-specific fluctuations, but rather reflected a reproducible property of the benchmark and of the proposed sequence representation. Taken together, the results summarized in Table 2 show that the proposed information-theoretic and compositional representation carries substantial discriminative signal, while also indicating that this signal is expressed predominantly in a local and nonlinear form. On this basis, KNN was selected as the main classifier for all subsequent binary-classification analyses. Importantly, the same qualitative ordering of models was also observed when the comparison was repeated on the original moderately imbalanced dataset, which comprised 212,962 sequences overall, including 81,481 fused and 131,481 non-fused samples. This confirms that the superiority of KNN was not merely an artifact of class balancing, but remained visible even when the full class prevalence structure of the original dataset was retained.

### 4.2. Final held out performance of the selected KNN classifier

After model selection by nested cross validation, we evaluated the selected K nearest neighbors classifier on an independent held out train, validation, and test partition of the balanced benchmark dataset. The benchmark was obtained by random undersampling of the majority class, resulting in 81,481 fused and 81,481 non fused sequences, which were then partitioned into 110,813 training instances, 19,556 validation instances, and 32,593 test instances. The final classifier used the hyperparameter configuration that was most stable across nested cross validation, namely *k* = 5, distance based neighbor weighting, and Minkowski distance with *p* = 1.

Two decision thresholds were defined on the validation set, each serving a different purpose:

- thr_*high*_: selected by maximizing the validation F1 score and used as the main operating point for the standalone fused versus non fused classification task, where the goal is to balance precision and recall.
- thr_*low*_: selected as the highest threshold achieving validation recall of at least 0.90 and used as a more permissive screening threshold for the downstream two stage Pipeline Detection+Decoding, where retaining potentially fused candidates is prioritized over first stage precision.

In practical terms, these two thresholds define three operating regions for the first stage classifier. A sequence assigned a fusion probability below thr_*low*_ is discarded as insufficiently fusion like, whereas a sequence scoring above thr_*high*_ is retained even under the stricter standalone binary classification setting. By contrast, sequences with scores in the intermediate range between thr_*low*_ and thr_*high*_ are the borderline cases that are filtered differently depending on the analytical objective. For example, with thr_*low*_ = 0.51 and thr_*high*_ = 0.62, a sequence receiving a predicted fusion probability of 0.57 would be passed to the second stage partner gene and breakpoint predictor, but it would not be called fused in the main binary classification setting. In this way, thr_*low*_ functions as a sensitivity oriented gate that protects candidate fused sequences from being discarded too early, whereas thr_*high*_ defines the stricter operating point used when a direct fused versus non fused decision is required.

As expected, these two thresholds produced different threshold dependent metrics, whereas AUROC and AUPRC remained unchanged because they do not depend on a fixed decision threshold. At the main classification operating point, thr_*high*_ = 0.62, the final KNN classifier achieved an AUROC of 0.892 with a 95% bootstrap confidence interval of [0.888, 0.895] and an AUPRC of 0.865 with a 95% bootstrap confidence interval of [0.860, 0.871] on the held out test set. Threshold dependent performance was also strong, with an accuracy of 0.834 [0.831, 0.838], an F1 score of 0.835 [0.831, 0.840], a precision of 0.831 [0.826, 0.837], and a recall of 0.840 [0.834, 0.845] (Table 3). When the more permissive screening threshold was used, thr_*low*_ = 0.51, recall increased to 0.899 [0.895, 0.904], while precision decreased to 0.741 [0.736, 0.747], with corresponding reductions in accuracy and F1 score. This behavior is entirely consistent with the intended role of thr_*low*_, since the lower threshold admits a broader set of candidate fused sequences into the downstream pipeline and therefore sacrifices first stage precision in exchange for higher sensitivity. A direct comparison between the two thresholds is reported in Table 4.

**Table 3.** Final held out performance of the selected KNN classifier on the balanced benchmark dataset at the main binary classification operating point, defined by the validation selected F1 maximizing threshold thr_*high*_ = 0.62. Confidence intervals were estimated by bootstrap resampling of the held out test set.

| Metric | KNN ($\text{thr}_{\text{high}} = 0.62$) |
| --- | --- |
| AUROC | 0.892 [0.888, 0.895] |
| AUPRC | 0.865 [0.860, 0.871] |
| Accuracy | 0.834 [0.831, 0.838] |
| F1 score | 0.835 [0.831, 0.840] |
| Precision | 0.831 [0.826, 0.837] |
| Recall | 0.840 [0.834, 0.845] |

**Table 4.** Comparison of the two operating thresholds used for the final held-out KNN classifier on the balanced benchmark dataset. The high threshold (thr_*high*_) was selected by maximizing validation F1-score, whereas the low threshold (thr_*low*_) was selected as the highest threshold achieving validation recall of at least 0.90. Confidence intervals were estimated by bootstrap resampling of the held-out test set.

| Metric | $\text{thr}_{\text{high}} = 0.62$ | $\text{thr}_{\text{low}} = 0.51$ |
| --- | --- | --- |
| Accuracy | 0.834 [0.831, 0.838] | 0.793 [0.788, 0.797] |
| F1-score | 0.835 [0.831, 0.840] | 0.813 [0.808, 0.817] |
| Precision | 0.831 [0.826, 0.837] | 0.741 [0.736, 0.747] |
| Recall | 0.840 [0.834, 0.845] | 0.899 [0.895, 0.904] |

Overall, these held out results confirm that the KNN classifier selected by nested cross validation generalizes well beyond the model selection procedure and preserves strong discriminative performance on unseen data. At the same time, they show that the same classifier can be operated in two distinct regimes, one optimized for balanced binary classification and one optimized for sensitive candidate selection prior to downstream fusion characterization.

### 4.3. Resolved mutual-information descriptors are the main source of discriminative signal

To determine which components of the proposed representation were most responsible for fused-versus-non-fused discrimination, we examined the final 31-feature space through two complementary analyses, namely permutation importance at the individual-feature level and feature-family ablation at the group level. Although these two approaches probe the representation from different angles, they converged on the same conclusion and consistently identified nucleotide-resolved mutual-information descriptors as the dominant source of predictive signal.

At the level of individual variables, permutation importance on the validation set showed a clear predominance of resolved mutual-information features among the highest-ranked descriptors, with the strongest performance decreases observed when variables at the shortest lag (*d* = 1) were permuted. This indicates that very local nucleotide-dependency patterns carry the most informative signal for binary classification, while additional resolved mutual-information terms at lags *d* = 2 and *d* = 3 also remained highly ranked and therefore point to a broader contribution of short-range sequence organization. By comparison, among the non-mutual-information features only GC content appeared consistently near the top of the ranking, and even in that case its contribution remained smaller than that of the leading resolved mutual-information variables.

The same picture emerged even more clearly from the feature-family ablation analysis summarized in Table 5. When the full 31-feature representation was used, the KNN classifier reached an AUROC of 0.894 and an AUPRC of 0.867, whereas removing the resolved mutual-information family caused a substantial deterioration in performance, reducing AUROC to 0.794 and AUPRC to 0.771, together with marked losses in accuracy and F1-score. Conversely, a model trained using only resolved mutual-information descriptors still achieved an AUROC of 0.892 and an AUPRC of 0.867, thus coming very close to the performance of the complete feature space. This near equivalence between the full model and the rMI-only model indicates that most of the predictive structure of the benchmark is already concentrated in short-range nucleotide dependency patterns.

**Table 5.** Feature-family ablation analysis for the KNN classifier on the final 31-feature representation. Performance is reported on the held-out split using the validation-selected decision threshold.

| Setting | AUROC | AUPRC | Accuracy | F1-score |
| --- | --- | --- | --- | --- |
| All features | 0.894 | 0.867 | 0.837 | 0.839 |
| Drop entropy | 0.897 | 0.872 | 0.841 | 0.843 |
| Drop GC | 0.894 | 0.866 | 0.838 | 0.840 |
| Only rMI | 0.892 | 0.867 | 0.835 | 0.837 |
| Drop rMI | 0.794 | 0.771 | 0.696 | 0.742 |
| Only entropy | 0.515 | 0.513 | 0.505 | 0.660 |
| Only GC | 0.511 | 0.503 | 0.514 | 0.646 |

Global compositional summaries, in contrast, were far less informative when considered in isolation. The entropy-only model reached an AUROC of 0.515 and the GC-only model an AUROC of 0.511, both values lying close to random expectation, while removing entropy-based descriptors from the full model slightly improved performance and removing GC content had only a negligible effect. Taken together, these results show that fused-versus-non-fused discrimination is driven much more strongly by local dependency structure than by global compositional biases alone. Overall, the close agreement between permutation importance and feature-family ablation provides a coherent interpretation of the benchmark and supports the central idea of this study, namely that fusion-associated sequences are characterized not only by broad compositional properties but, more importantly, by distinctive short-range dependency patterns that can be captured effectively through interpretable information-theoretic descriptors. Notably, the near equivalence between the full model and the model using only resolved mutual-information descriptors indicates that most of the predictive structure of the benchmark is already concentrated in short-range nucleotide dependency patterns.

### 4.4. Fusion partner and breakpoint prediction on fused sequences

We next evaluated the second-stage multitask model on the fused-only test set under the *Oracle* setting, that is, under the idealized assumption that the input sequences were already known to belong to the positive class. This analysis is important because it isolates the intrinsic capacity of the second-stage model from any error introduced by the first-stage fused-versus-non-fused filter and therefore provides a direct estimate of how much partner-gene and breakpoint information is recoverable from the same 31-feature sequence representation once fusion status is no longer uncertain. The Oracle results are reported together with the two-stage Detection+Decoding results in Table 6.

**Table 6.** Second-stage multitask performance under Oracle and two-stage Detection+Decoding evaluation settings. In the Oracle setting, the Decoding model was evaluated directly on the fused-only test set. In the two-stage setting, Decoding was applied only to sequences classified as fusion-positive by the Detection stage, using either the permissive threshold thr_*low*_ or the stricter threshold thr_*high*_. Partner-gene prediction is reported as top-*k* accuracy, whereas breakpoint metrics are reported on the normalized breakpoint coordinate.

| Metric | Oracle | Detection+Decoding $\text{thr}_{\text{low}}$ | Detection+Decoding $\text{thr}_{\text{high}}$ |
| --- | --- | --- | --- |
| Gene1 top-1 accuracy | 0.727 | 0.784 | 0.829 |
| Gene1 top-3 accuracy | 0.806 | 0.862 | 0.902 |
| Gene1 top-5 accuracy | 0.829 | 0.882 | 0.920 |
| Gene2 top-1 accuracy | 0.819 | 0.874 | 0.911 |
| Gene2 top-3 accuracy | 0.873 | 0.923 | 0.954 |
| Gene2 top-5 accuracy | 0.888 | 0.935 | 0.963 |
| Breakpoint MAE | 0.179 | 0.179 | 0.180 |
| Breakpoint MSE | 0.048 | 0.048 | 0.048 |

Under this setting, the model showed strong and highly structured performance for both partner-gene prediction tasks, while breakpoint prediction remained stable across the fused test set. For *gene1*, top-1 accuracy reached 0.727 (95% CI [0.721, 0.734]), increased to 0.806 (95% CI [0.801, 0.812]) at top-3, and further rose to 0.829 (95% CI [0.823, 0.834]) at top-5. Performance was consistently higher for *gene2*, for which the model reached 0.819 (95% CI [0.813, 0.825]) at top-1, 0.873 (95% CI [0.868, 0.878]) at top-3, and 0.888 (95% CI [0.882, 0.892]) at top-5. The systematic improvement from top-1 to top-5 in both tasks indicates that the correct partner label is often placed among the most plausible candidates even when it is not ranked first, whereas the consistently stronger performance observed for *gene2* suggests that the second partner label, as represented in the present dataset, is more directly recoverable from the extracted descriptors than *gene1*.

Breakpoint prediction was more moderate in absolute terms but remained remarkably stable, with a mean absolute error of 0.179 (95% CI [0.177, 0.181]) and a mean squared error of 0.048 (95% CI [0.047, 0.049]) on the normalized breakpoint coordinate. This indicates that the information-theoretic and compositional descriptors retain usable signal not only for partner-gene identification but also for approximate breakpoint localization, although the latter appears to be encoded in a more distributed and less sharply classifiable form than the partner labels.

Taken together, the Oracle results in Table 6 show that the proposed sequence representation supports a substantially richer characterization of fusion-positive sequences than binary classification alone. Once the model is restricted to authentic fused sequences, the same descriptor space retains enough structure to recover both partner-gene identity and coarse breakpoint position. This, in turn, motivates the next question addressed by the pipeline, namely how much of this second-stage performance is preserved when the multitask model is applied not in the Oracle setting, but only after candidate sequences have first been screened by the initial KNN classifier.

### 4.5. The two-stage Pipeline Detection+Decoding markedly enriches downstream partner prediction

We then moved from the idealized Oracle setting to the full two-stage configuration, in which the multitask model of Pipeline Decoding was applied only to sequences that had first been classified as fusion-positive by the KNN classifier of Pipeline Detection. In the updated evaluation scheme, both stages inherited the same master train, validation, and test partition, so that the Oracle and Pipeline Detection+Decoding results were directly comparable on aligned data splits. This point is important, because it allows the downstream gains observed in Table 6 to be interpreted as a genuine enrichment effect introduced by first-stage screening rather than as a consequence of mismatched partitioning.

Under the Oracle setting, in which Pipeline Decoding was evaluated on the full fused-only test subset, partner prediction was already strong, with *gene1* top-1, top-3, and top-5 accuracies of 0.727, 0.806, and 0.829, respectively, and corresponding *gene2* accuracies of 0.819, 0.873, and 0.888. However, once Pipeline Decoding was restricted to the sequences passing the first-stage KNN filter, performance increased sharply and consistently across all top-*k* metrics. This enrichment effect was already evident under the more permissive threshold thr_*low*_, which was selected as the highest validation threshold achieving recall of at least 0.90 in Pipeline Detection. In this setting, *gene1* top-1, top-3, and top-5 accuracy increased from 0.727, 0.806, and 0.829 to 0.784, 0.862, and 0.882, while *gene2* improved from 0.819, 0.873, and 0.888 to 0.874, 0.923, and 0.935. Thus, even a sensitivity-oriented first-stage gate enriched the candidate set for fusion-positive sequences whose partner architecture was much easier to recover from the same information-theoretic representation. Importantly, bootstrap confidence intervals for the thr_*low*_ setting remained narrow, with *gene1* top-1, top-3, and top-5 accuracies of 0.784 [0.778, 0.791], 0.862 [0.856, 0.868], and 0.882 [0.877, 0.887], and corresponding *gene2* accuracies of 0.874 [0.869, 0.880], 0.923 [0.919, 0.928], and 0.935 [0.931, 0.939], confirming that this gain was stable and not driven by random fluctuations of the test set. The effect became even stronger under the stricter threshold thr_*high*_, which had been selected by maximizing the validation F1-score of Pipeline Detection and therefore identifies a more confident subset of fusion-like sequences. In that regime, *gene1* top-1, top-3, and top-5 accuracy rose further to 0.829, 0.902, and 0.920, whereas *gene2* reached 0.911, 0.954, and 0.963. The improvement was therefore not marginal, but large and highly consistent across both partner labels and across all top-*k* metrics, showing that progressively stricter first-stage gating produces progressively more informative downstream candidates. By contrast, breakpoint regression remained essentially unchanged across the three evaluation settings. The normalized breakpoint MAE was 0.179 in the Oracle setting, 0.179 under thr_*low*_, and 0.180 under thr_*high*_, while MSE remained stable at approximately 0.048 throughout. The corresponding bootstrap intervals were also highly similar, with Oracle MAE and MSE equal to 0.179 [0.177, 0.181] and 0.048 [0.047, 0.049], thr_*low*_ MAE and MSE equal to 0.179 [0.177, 0.181] and 0.048 [0.047, 0.049], and thr_*high*_ MAE and MSE equal to 0.180 [0.178, 0.182] and 0.048 [0.047, 0.049].

This stability indicates that the main contribution of first-stage filtering concerns partner-gene recoverability rather than breakpoint localization, which appears to depend on a component of the sequence signal that is much less affected by candidate selection.

Taken together, these results show that the first-stage KNN classifier acts as more than a binary detector. By screening the candidate space before the multitask model is applied, Pipeline Detection preferentially forwards those fusion-positive instances whose partner architecture is also more readily recoverable, thereby functioning as an effective enrichment layer for downstream fusion characterization rather than as a simple pre-filter.

### 4.6. Robustness under fusion-pair-disjoint evaluation

To assess whether the predictive signal captured by the proposed sequence descriptors extends beyond previously observed fusion identities, we repeated the fused *versus* non-fused classification task under a substantially stricter grouped evaluation protocol. In this analysis, the balanced 31-feature dataset was first constructed by retaining all 81,481 fused samples and randomly undersampling the negative class to the same size. Positive instances were then grouped by fusion-pair identity, defined as the ordered *gene1*–*gene2* combination, whereas negative samples were assigned unique singleton groups. Repeated train–test partitions were generated with `GroupShuffleSplit`, using 10 independent repeats and a test fraction of 0.20, while enforcing zero overlap in positive fusion-pair identity between training and test sets. KNN and the feed-forward neural network were then evaluated on the same grouped splits, so that performance differences reflected model behavior rather than differences in partitioning (Table 7).

**Table 7.** Performance across auxiliary evaluation protocols. The FusionAI-like benchmark consisted of a single 70/30 train–test split on 26,000 real fused sequences and 26,000 synthetic pseudo-fusions generated from Ensembl cDNA reference transcripts. The fusion-pair-disjoint evaluation was performed on the balanced 31-feature dataset using 10 repeated `GroupShuffleSplit` partitions, with positive samples grouped by ordered fusion-pair identity and negative samples assigned unique singleton groups. FusionAI-like values are point estimates from the held-out test set, whereas fusion-pair-disjoint values are mean scores with approximate 95% confidence intervals.

| Metric | FusionAI-like KNN | Fusion-pair-disjoint KNN | Fusion-pair-disjoint FFNN |
| --- | --- | --- | --- |
| AUROC | 0.832 | 0.715 [0.712, 0.718] | 0.706 [0.703, 0.709] |
| AUPRC | 0.800 | 0.686 [0.680, 0.693] | 0.692 [0.685, 0.700] |
| Accuracy | 0.730 | 0.653 [0.650, 0.656] | 0.639 [0.636, 0.642] |
| F1-score | 0.764 | 0.632 [0.628, 0.636] | 0.618 [0.610, 0.626] |
| Precision | 0.678 | 0.674 [0.668, 0.680] | 0.659 [0.652, 0.665] |
| Recall | 0.875 | 0.595 [0.589, 0.601] | 0.582 [0.571, 0.594] |

As expected, performance decreased relative to the main random-split benchmark, confirming that prediction on previously unseen fusion pairs is a considerably harder task. Under this protocol, KNN achieved a mean AUROC of 0.715 (95% CI [0.712, 0.718]) and a mean AUPRC of 0.686 (95% CI [0.680, 0.693]), together with a mean accuracy of 0.653 (95% CI [0.650, 0.656]) and a mean F1-score of 0.632 (95% CI [0.628, 0.636]). The feed-forward neural network reached a mean AUROC of 0.706 (95% CI [0.703, 0.709]) and a mean AUPRC of 0.692 (95% CI [0.685, 0.700]), with a mean accuracy of 0.639 (95% CI [0.636, 0.642]) and a mean F1-score of 0.618 (95% CI [0.610, 0.626]).

Although this evaluation setting is intentionally severe, both models remained clearly above random expectation, indicating that the information-theoretic representation retains transferable predictive signal even when the classifier is forced to operate on fusion pairs that were never encountered during training. KNN remained the stronger model for AUROC, accuracy, F1-score, precision, and recall, whereas the neural network achieved a slightly higher mean AUPRC. This pattern suggests that local neighborhood structure still provides the most robust overall discrimination under stressed generalization, even though the neural model preserves a competitive ranking ability in terms of precision–recall behavior.

Overall, these results show that the proposed feature space does not merely capture identity-specific regularities of known fusion pairs, but also preserves a measurable capacity to generalize under a deliberately more demanding regime in which the model must extrapolate to previously unseen fusion architectures.

**<H12>4.7. Comparison with FusionAI-like protocols**

To further contextualize the proposed framework against a more explicitly breakpoint-centered sequence-classification scenario, we designed an auxiliary FusionAI-like evaluation protocol that attempted, as closely as possible within our setting, to mimic the logic of a pseudo-fusion benchmark. This comparison should be understood as a deliberate approximation rather than as a strict reproduction of the original FusionAI framework, because the two approaches are built on different assumptions: FusionAI is centered on breakpoint-proximal sequence modeling, whereas our method is based on compact alignment-free information-theoretic and compositional descriptors computed over the full sequence representation. Nevertheless, we sought to construct a comparison protocol that would be as similar as possible in spirit, so as to test whether the same descriptor space retained discriminative value when the negative class was no longer composed of matched non-fused controls, but instead of artificially generated pseudo-fusions. To this end, we assembled a balanced 1:1 benchmark containing 26,000 real fusion-positive instances and 26,000 synthetic pseudo-fusion negatives. Positive samples were randomly drawn from the positive class of the main dataset, whereas pseudo-fusion negatives were generated from transcript-derived reference sequences obtained from Ensembl cDNA annotations (release 115, GRCh38). For each gene, the longest available transcript was retained as the sequence source, after which candidate genes used to build pseudo-fusions were restricted to genes not observed among the fusion partners present in the sampled positive subset. Synthetic negatives were then created by joining transcript fragments from these non-partner genes, while matching the final sequence lengths to those of the sampled positive instances in order to reduce trivial length-driven effects. Once the auxiliary dataset had been generated, the same 31 information-theoretic and compositional descriptors used throughout the main study were recomputed for all samples, so that the comparison differed in benchmark construction rather than in feature representation.

The resulting FusionAI-like dataset was evaluated by means of a 70/30 train-test split, and under this protocol the K-nearest-neighbors classifier achieved an AUROC of 0.832, an AUPRC of 0.800, an accuracy of 0.730, a precision of 0.678, a recall of 0.875, and an F1-score of 0.764 (Table 7). Although these values are lower than those obtained on the main benchmark, they remain substantial and indicate that the proposed descriptor space preserves meaningful discriminative signal even when the negative class is defined by synthetic chimeric constructs rather than by naturally occurring non-fused controls.

This result is relevant for two reasons. First, it shows that the predictive information captured by entropy-based, mutual-information-based, and compositional descriptors is not confined to a single benchmark design, but remains detectable under a substantially altered negative-class construction. Second, it highlights the conceptual difference between the present framework and FusionAI-like paradigms: in our case, the method is not intrinsically designed around pseudo-fusion generation, and therefore this experiment constitutes, by construction, a methodological forcing of our representation into a different evaluation regime. The fact that performance remains robust under this forcing provides additional support for the generality of the proposed sequence representation.

### 4.8. Comparison with alignment based fusion callers

To position the proposed alignment free framework with respect to established fusion detection pipelines, we also considered two widely used alignment based callers, namely *STAR-Fusion* and *FusionCatcher*. The comparison was based on their ability to recover the correct fusion partner genes from the same underlying dataset used in this study. Because these tools are designed for fusion event discovery rather than for direct binary classification of pre extracted sequence instances, their outputs required an auxiliary evaluation procedure in order to be mapped onto our benchmark. In practice, the alignment based callers often produced ambiguous outputs for the same case, including multiple candidate fusion events or uncertain partner assignments. This behavior reflects the nature of event discovery pipelines, which are designed to report plausible fusion hypotheses from alignment evidence rather than to return a single unambiguous decision for each sequence instance. For this reason, the derived evaluation protocol labelled a read as positive if it contributed to a reported fusion event associated with the expected fusion partners, and negative otherwise.

Metrics for *STAR-Fusion* and *FusionCatcher* were therefore computed at read level, whereas the proposed framework was evaluated directly at sequence level on a held out benchmark.

Within this contextual evaluation setting, *STAR-Fusion* achieved an accuracy of 0.60, a ROC AUC of 0.60, a precision of 0.99, a recall of 0.60, and an F1 score of 0.75 (Table 8). This pattern is consistent with a highly conservative calling behavior, characterized by very high precision but only moderate recall. By contrast, *FusionCatcher* achieved an accuracy of 0.27, a ROC AUC of 0.60, a precision of 0.28, a recall of 0.60, and an F1 score of 0.53, indicating substantially greater ambiguity and lower selectivity under the same evaluation scheme.

**Table 8.** Contextual comparison between the proposed method and two widely used alignment based fusion callers. *STAR-Fusion* and *FusionCatcher* were evaluated according to their ability to recover the expected fusion partner genes under a derived read level protocol. In this setting, ambiguous or multiple candidate calls could be produced for the same case. The proposed method is reported for the held out KNN classifier on the main sequence level benchmark and is shown in bold for reference.

| Method | Accuracy | ROC AUC | Precision | Recall | F1 score |
| --- | --- | --- | --- | --- | --- |
| STAR-Fusion | 0.60 | 0.60 | 0.99 | 0.60 | 0.75 |
| FusionCatcher | 0.27 | 0.60 | 0.28 | 0.60 | 0.53 |
| Proposed method | **0.834** | **0.892** | **0.831** | **0.840** | **0.835** |

These values should therefore be interpreted as contextual rather than as a strictly matched one to one benchmark. The purpose of this analysis was not to claim full methodological equivalence between event level fusion discovery and instance level sequence classification, but to place the proposed framework relative to widely used fusion analysis paradigms. In this perspective, the comparison highlights that fusion related signal can be approached not only through alignment, split read support, and event reconstruction, but also through intrinsic sequence organization captured by entropy and mutual information derived descriptors. This supports the view that the two paradigms are complementary rather than interchangeable.

### 4.9. Internal breakpoint centered validation

To determine whether the descriptors highlighted by the predictive analyses also captured biologically meaningful local properties of fusion regions, we performed two complementary breakpoint-centered internal validations. These analyses were important because the main classification models were trained on global sequence-level representations, and therefore strong predictive performance alone would not be sufficient to show that the retained signal was actually related to local breakpoint biology rather than to more diffuse or potentially artifactual statistical differences across classes. For this reason, we asked whether the same feature families that proved informative at the global level also showed coherent behavior in the immediate sequence context of authentic fusion breakpoints.

More specifically, we carried out: (I) a local window-based validation, in which fixed-length windows centered on real fusion breakpoints were compared with matched windows extracted from non-fused control sequences at the same relative position, after recomputing local information-theoretic descriptors within each window; and (II) a breakpoint microhomology analysis, in which exact junction microhomology was quantified and compared between fusion-positive sequences and matched non-fusion controls. Taken together, these two analyses were designed to test whether the feature space emphasized by the predictive models also reflected plausible local sequence properties associated with fusion formation and rearrangement-related repair.

In the local window analysis, the strongest univariate differences were observed for entropy based descriptors. Breakpoint centered fusion windows showed higher local Shannon entropy and higher local Rényi entropy than matched non fused control windows, with absolute univariate AUROC values of 0.608 for both descriptors (Table 9; Figure 1). These results indicate that true fusion breakpoint regions tend to occur in locally more heterogeneous sequence contexts. Short range dependency descriptors also showed consistent, although smaller, effects. The most informative mutual information based signals were concentrated at lag 1, including both the total mutual information at lag 1 and nucleotide resolved components, especially the cytosine resolved contribution. In general, fusion centered windows were characterized by altered immediate nucleotide dependency structure relative to matched non fused windows, whereas effects at longer lags were weaker.

**Table 9.** Top breakpoint-centered local features discriminating real fusion windows from matched non-fused control windows. AUROC values are reported as absolute univariate discrimination performance.

| Feature | AUROC | Cliff's $\delta$ | Direction in fusion windows |
| --- | --- | --- | --- |
| Local Shannon entropy | 0.608 | 0.217 | higher |
| Local Rényi entropy ($\alpha = 2$) | 0.608 | 0.216 | higher |
| Local C-resolved mutual information ($\tau = 1$) | 0.563 | -0.126 | lower |
| Local mutual information ($\tau = 1$) | 0.543 | -0.086 | lower |
| Local A-resolved mutual information ($\tau = 1$) | 0.531 | -0.061 | lower |
| Local T-resolved mutual information ($\tau = 1$) | 0.526 | 0.051 | higher |

**Figure 1.**
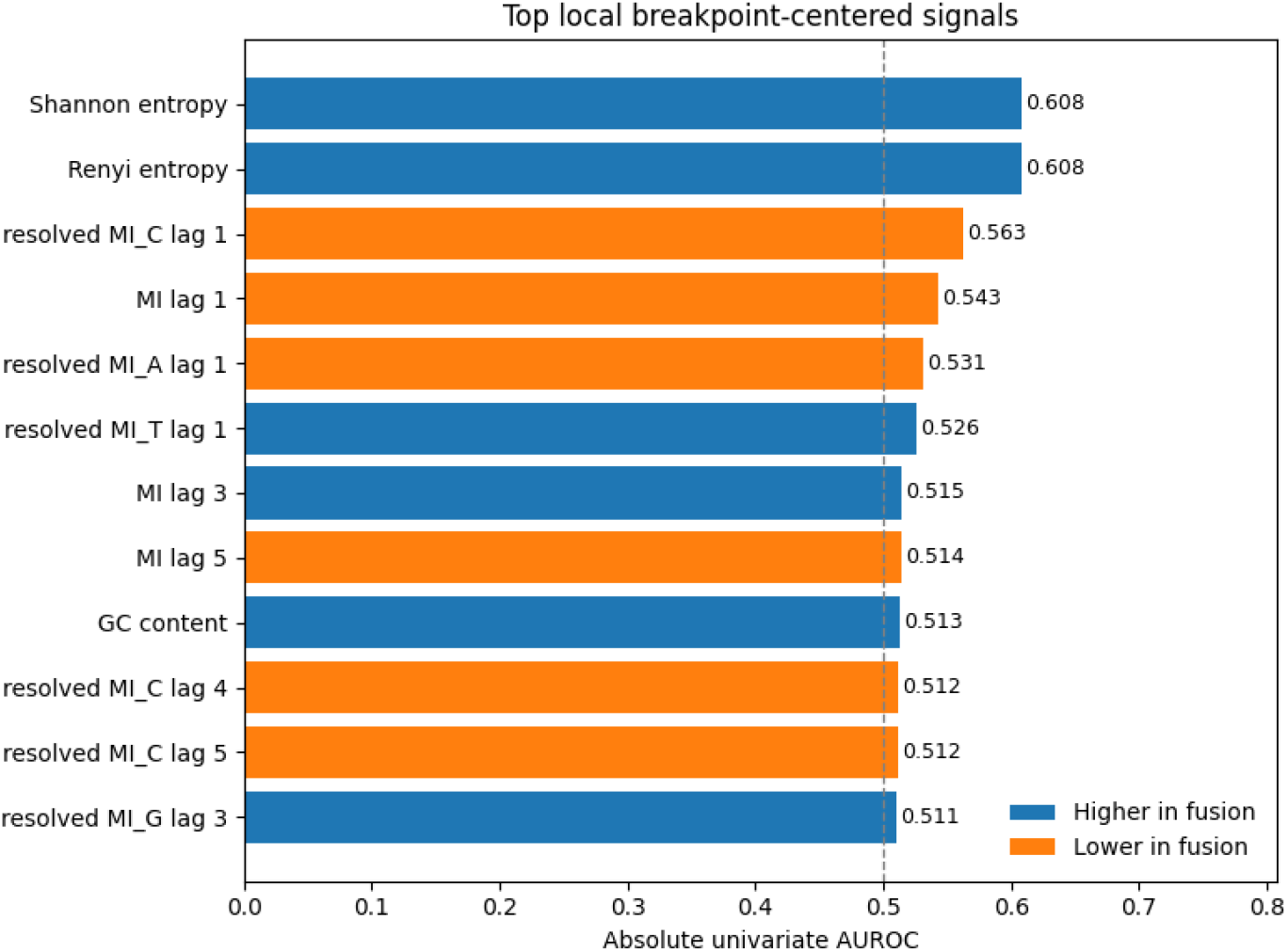
Top local breakpoint centered descriptors discriminating real fusion windows from matched non fused control windows. Bars are coloured according to whether the corresponding descriptor is higher or lower in fusion windows. The strongest local signals were observed for entropy based descriptors and for short range mutual information based features, especially at lag 1. Absolute univariate AUROC values are shown on the horizontal axis, and the dashed line marks random expectation.

To visualize how these local signals varied around the breakpoint, we further summarized the descriptors as a function of distance from the junction (Figure 2).

**Figure 2.**
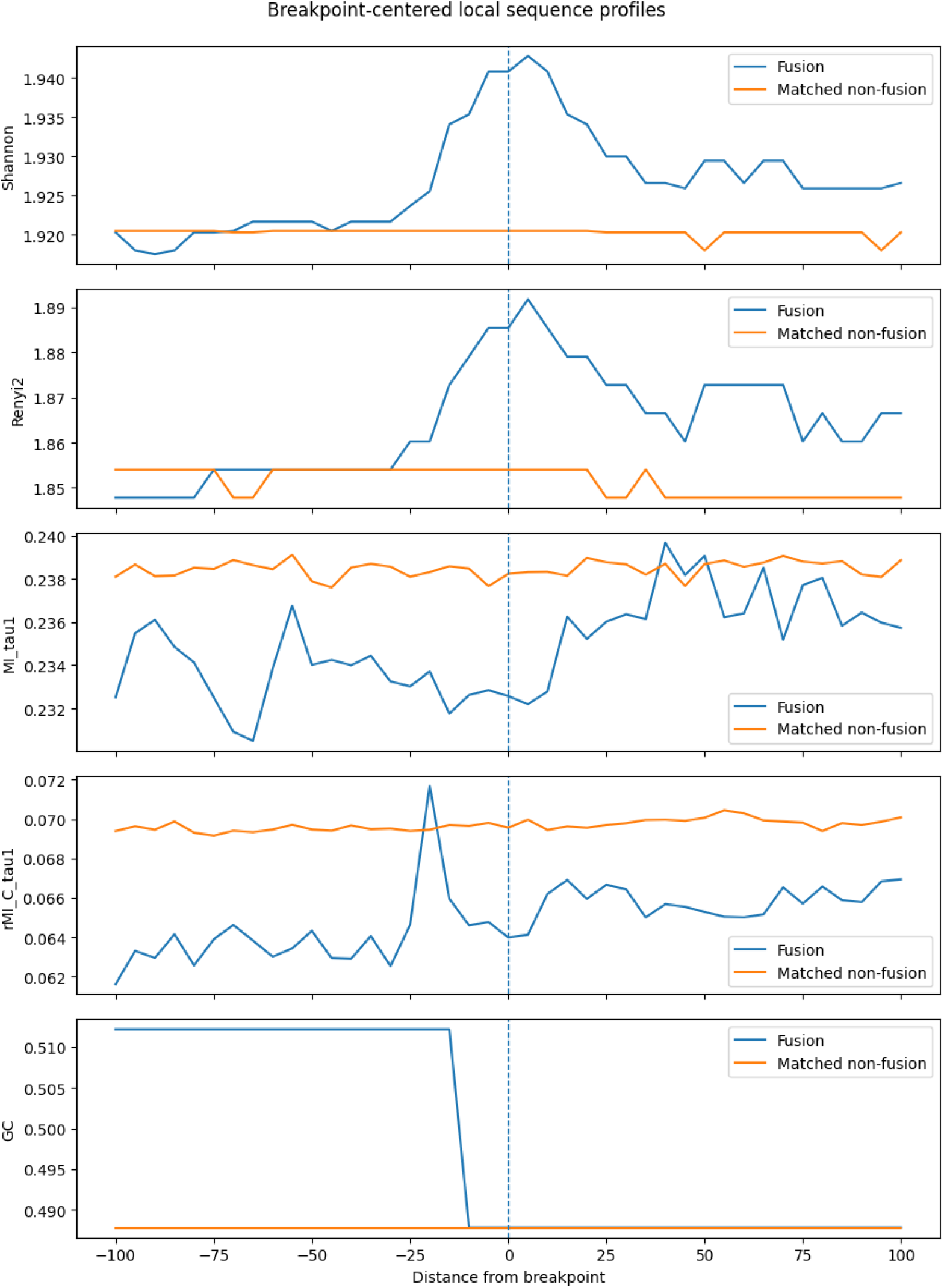
Breakpoint centered local sequence profiles in fusion and matched non fused windows. Curves show the median local descriptor value as a function of distance from the breakpoint. The vertical dashed line marks the breakpoint position. Entropy based descriptors increase near the breakpoint in fusion windows, whereas mutual information based descriptors reveal altered short range dependency structure relative to matched controls.

Entropy based measures showed a local elevation in fusion windows near the breakpoint, whereas mutual information based descriptors displayed altered short range dependency structure relative to controls. Although descriptive, these profiles supported the same overall interpretation emerging from the univariate ranking analysis. We next examined breakpoint microhomology as a complementary mechanistic validation of fusion associated local sequence structure. Using a conservative matched analysis, fusion positive samples showed a significant enrichment of exact breakpoint microhomology relative to non fusion controls (Table 10; Figure 3). Although the median microhomology length was 0 in both groups, fusion positive sequences displayed a higher mean microhomology length (0.570 vs. 0.513) and a larger fraction of non zero microhomology events (0.469 vs. 0.353). This difference was highly significant by Mann–Whitney testing (*p* = 1.76 × 10^−16^), although the effect size remained modest (Cliff’s *δ* = 0.091). Consistently, microhomology alone had limited discriminative power, with a univariate AUROC of 0.545.

**Table 10.**
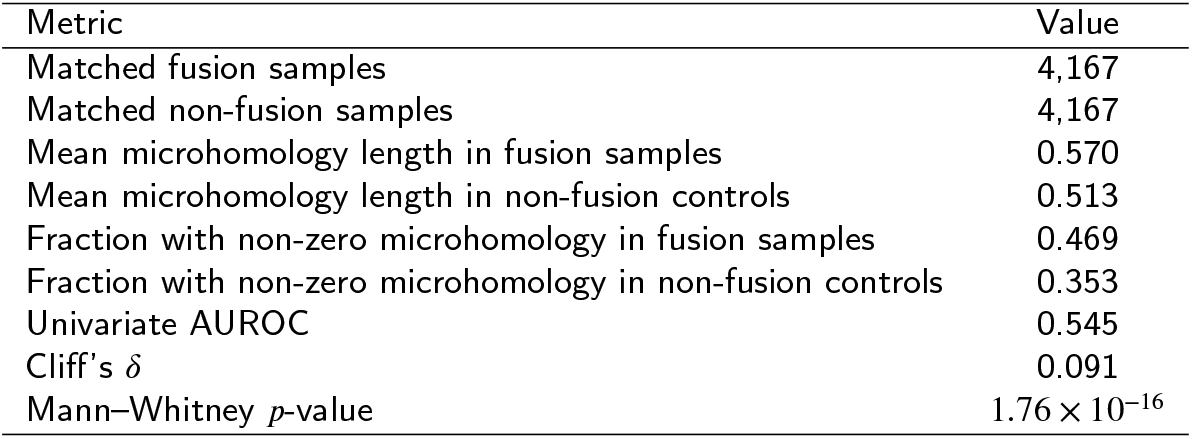
Breakpoint microhomology enrichment in fusion-positive sequences relative to matched non-fusion controls.

| Metric | Value |
| --- | --- |
| Matched fusion samples | 4,167 |
| Matched non-fusion samples | 4,167 |
| Mean microhomology length in fusion samples | 0.570 |
| Mean microhomology length in non-fusion controls | 0.513 |
| Fraction with non-zero microhomology in fusion samples | 0.469 |
| Fraction with non-zero microhomology in non-fusion controls | 0.353 |
| Univariate AUROC | 0.545 |
| Cliff's $\delta$ | 0.091 |
| Mann–Whitney $p$-value | $1.76 \times 10^{-16}$ |

**Figure 3.**
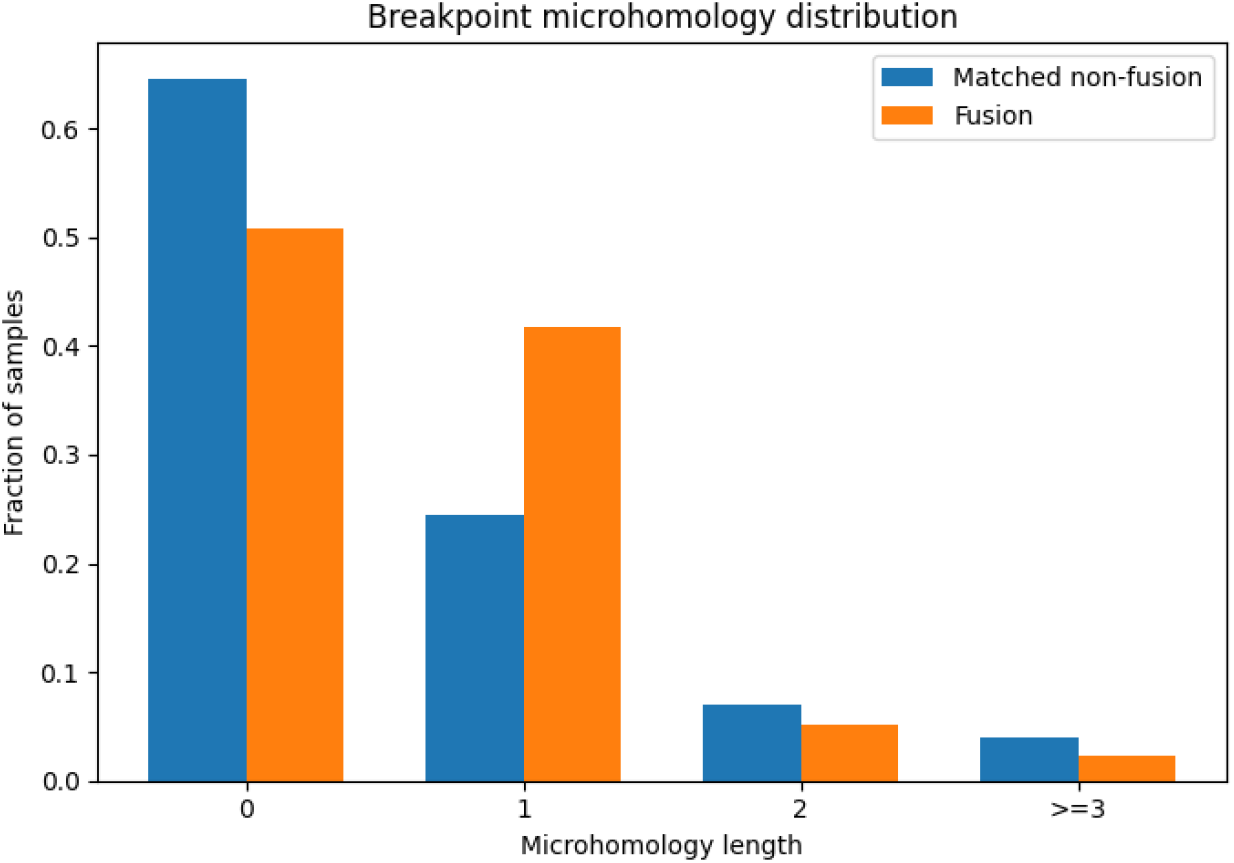
Distribution of breakpoint microhomology length in matched fusion and non fused windows. Fusion positive regions show a reduced fraction of zero microhomology and a corresponding enrichment of short non zero microhomology, consistent with the idea that authentic fusion breakpoint regions are modestly enriched in sequence contexts compatible with rearrangement associated repair processes.

The relationship between breakpoint microhomology and the global handcrafted features was generally weak, suggesting that microhomology explains only a limited fraction of the signal captured by the classifier. Small associations were observed for GC content and selected mutual information descriptors, but no strong monotonic dependence emerged. This indicates that the predictive value of the feature space does not reduce to breakpoint overlap alone. Taken together, these internal validations support a coherent biological interpretation of the proposed representation. Fusion associated regions appear to be characterized by increased local sequence complexity, altered short range nucleotide dependency structure, and a modest enrichment of breakpoint microhomology. The local entropy and mutual information results are consistent with the feature importance and ablation analyses, which identified mutual information based descriptors as dominant contributors to binary classification, while the microhomology analysis provides an additional mechanistic link between fusion breakpoints and local sequence architecture. Overall, these findings suggest that the handcrafted descriptors capture biologically meaningful sequence organization rather than merely providing abstract statistical separation.

Although the median microhomology length was 0 in both groups, the distribution of breakpoint microhomology was shifted toward higher values in fusion-positive sequences, which also showed a larger fraction of non-zero events and a higher mean microhomology length (Fig. 4).

**Figure 4.**
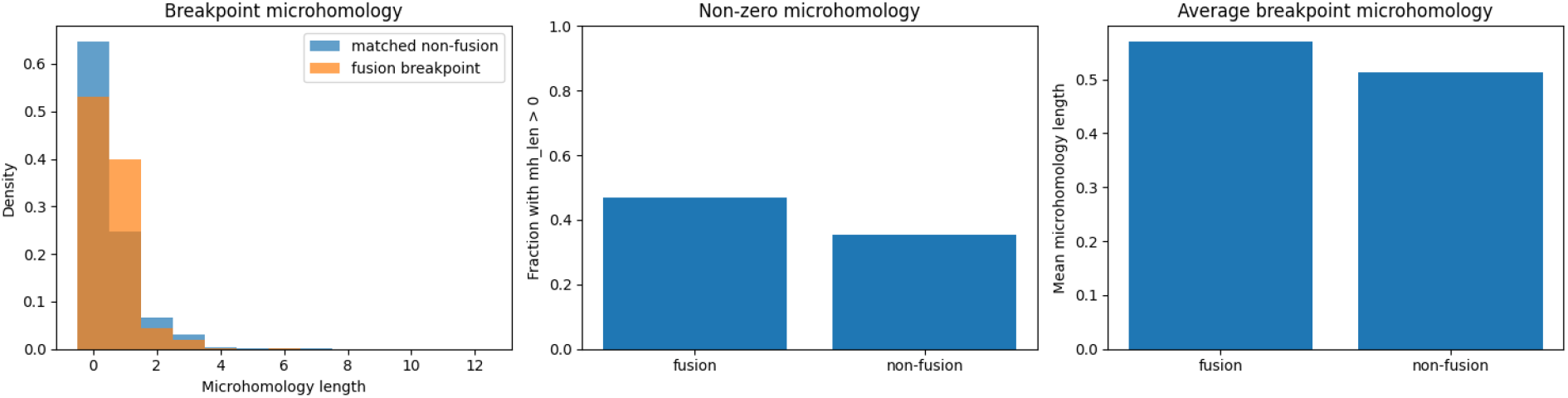
Breakpoint microhomology in fusion-positive sequences and matched non-fusion controls. Left: distribution of exact breakpoint microhomology length. Middle: fraction of samples with non-zero microhomology. Right: mean breakpoint microhomology length. Fusion-positive samples show a modest but consistent enrichment of breakpoint microhomology relative to matched non-fusion controls.

## 5. Discussion

The present study shows that a compact set of interpretable information theoretic and compositional descriptors captures substantial sequence level signal associated with gene fusions. Across the main benchmark, this representation supported accurate fused versus non fused classification, with K nearest neighbors consistently outperforming linear baselines and a feed forward neural network. This result is relevant not only from a predictive perspective, but also from a representational one, because it indicates that fusion related and non fusion related sequences can be separated using only intrinsic sequence organization, without relying on alignment, transcript reconstruction, or explicit breakpoint modelling.

The most important result of the study is that the predictive content of the feature space is not distributed uniformly across descriptors. Instead, resolved mutual information consistently emerged as the dominant source of signal. Both permutation importance and feature family ablation converged on the same conclusion: performance remained close to that of the full model when only resolved mutual information descriptors were retained, whereas it decreased sharply when this family was removed. By contrast, entropy and GC content alone showed limited predictive value. This recurring pattern strongly suggests that short range nucleotide dependency structure is one of the most informative statistical signatures of fusion associated sequences in this benchmark.

The internal validation analyses reinforce this interpretation. Breakpoint centered local windows showed higher local entropy and altered short range dependency structure relative to matched non fused controls, indicating that fusion regions are characterized by increased local sequence complexity together with modified nucleotide organization. In parallel, the microhomology analysis showed that fusion positive regions were enriched in non zero microhomology relative to matched controls, supporting the view that the detected sequence signal is linked to biologically meaningful breakpoint contexts. Although no single descriptor can fully explain this behavior on its own, the combined results suggest that the proposed representation captures a composite signature of local complexity, sequence dependency, and rearrangement associated context.

The downstream analyses further show that this handcrafted representation contains information beyond binary discrimination alone. On fused sequences, the second stage model achieved strong partner gene prediction together with stable breakpoint regression performance. The two stage Pipeline Detection+Decoding improved partner prediction after first stage filtering, while leaving breakpoint error essentially unchanged, indicating that the first classifier acts not only as a detector of fusion related signal but also as an enrichment step for downstream characterization.

The robustness experiments are also informative. Under fusion pair disjoint evaluation, performance decreased relative to the main random split benchmark, confirming that this is a much harder generalization task. Nevertheless, both KNN and the neural baseline remained clearly above random expectation, showing that the feature space does not merely memorize specific fusion identities and retains transferable signal even under stressed evaluation conditions. Additional contextual comparisons in FusionAI like and alignment based settings further support the idea that fusion related sequence information can be captured from a perspective that is complementary to alignment based event discovery.

Overall, the study indicates that fusion associated DNA segments contain detectable and interpretable information in their local statistical organization, and that this information can be captured effectively through a small number of handcrafted descriptors. Most importantly, the repeated emergence of resolved mutual information as the strongest component across classification, ablation, robustness, and internal validation analyses identifies short range nucleotide dependency structure as a central feature of fusion related sequence organization. In this sense, the present work does not simply provide an alignment free classifier, but also highlights a specific interpretable signal that may serve as a useful starting point for future biological investigation of fusion prone genomic contexts.

## CRediT authorship contribution statement

**Gerardo Benevento:**. **Delfina Malandrino:**. **Alessia Ture:**. **Rocco Zaccagnino:** .

## Notes

### Competing Interest Statement

The authors have declared no competing interest.

